# Seasonal evolution of *Drosophila melanogaster* abdominal pigmentation is associated with a multifarious selective landscape

**DOI:** 10.1101/2025.08.15.670575

**Authors:** Skyler Berardi, Jack K. Beltz, Seth M. Rudman, Tess N. Grainger, Jonathan M. Levine, Hayes Oken, Paul Schmidt

## Abstract

Pigmentation has been widely studied by evolutionary biologists due to both ease of measure and relationship to fitness. *Drosophila melanogaster* pigmentation has represented a particularly useful avenue of investigation, as extensive genetic tools have enabled the characterization of the trait’s complex architecture. *Drosophila* pigmentation also varies predictably across space and time in wild populations, suggesting pigmentation is a component of adaptation to local environmental conditions. Despite this, the impact of *D. melanogaster* pigmentation on fitness, and the environmental factors that drive the evolution of pigmentation, are not well understood. To address this gap, we experimentally evolved replicated *D. melanogaster* populations in field mesocosms to determine whether and how pigmentation evolves in response to environmental variation. We found that pigmentation rapidly and predictably adapted to a direct manipulation of temperature, supportive of melanization playing a role in thermoregulation. However, we also determined that pigmentation responded adaptively to direct manipulations of numerous additional factors, including intraspecific competition, diet, and the microbiome. These findings suggest that the selective landscape acting on pigmentation is complex and multifaceted, and that patterns of melanization may be driven, at least in part, by indirect selection due to correlations with other fitness-related traits.

## Introduction

Pigmentation represents an extensively studied phenotype across a breadth of taxa, and it is a particularly useful model for defining how selective pressures shape dynamics of adaptation. Patterns of coloration can be characterized visually, and links between genotype and phenotype have been established across wild populations of numerous species (Hoekstra, 2006; Kronforst et al., 2012). The adaptive significance of pigmentation is straightforward in many systems, which yields clear predictions for how environmental change will interact with patterns of coloration. Adaptive shifts in pigmentation are often driven by crypsis, with notable examples including melanism in the peppered moth (Cook et al., 2012) and the pygmy grasshopper (Forsman et al., 2011), blanched coloration in White Sands lizards (Rosenblum et al., 2004), coat color shifts in mice (Barrett et al., 2019), and seasonal camouflage in snowshoe hares (Jones et al., 2018; Zimova et al., 2016). Alternatively, pigmentation patterns are known to be shaped by selective pressures such as thermoregulation (Kingsolver, 1987; Watt, 1968), UV tolerance (Jablonski & Chaplin, 2010), or aposematism (Mappes et al., 2005). However, in other species, the connection between pigmentation and fitness is less clear, and this includes the model fly *Drosophila melanogaster*.

Despite *D. melanogaster* pigmentation being an established model for studying the genetic basis of complex trait development (Massey & Wittkopp, 2016; Wittkopp et al., 2003), we have yet to unambiguously identify the agents of selection that shape variance in melanization observed in natural populations. *Drosophila* pigmentation exhibits patterns suggestive of adaptation in wild populations: parallel clines in melanization have repeatedly emerged across latitudinal and altitudinal gradients on multiple continents, in which flies are consistently darker at both higher latitudes and higher altitudes (Bastide et al., 2014; Berardi et al., 2025; Das, 1995; David et al., 1985; Dev et al., 2013; Munjal et al., 1997; Parkash et al., 2008; Pool & Aquadro, 2007; Rajpurohit et al., 2008a; Telonis-Scott et al., 2011). These clinal patterns are concordant with the thermal melanism hypothesis, which predicts that darker pigmentation will be favored under colder conditions (e.g., at higher latitudes and altitudes) as a mechanism for elevating body temperature, and lighter pigmentation will be favored in warmer environments to mitigate overheating (Clusella-Trullas et al., 2007; Gibert et al., 2000; Gibson & Falls, 1979; Goulson, 1994; Kingsolver & Wiernasz, 1991; Watt, 1969).

While it was previously thought that *D. melanogaster* would be too small to thermoregulate via pigmentation (Willmer & Unwin, 1981), a recent study found that darker cuticle pigmentation is associated with a higher body temperature when flies are exposed to light in the laboratory (Freoa et al., 2023). Additionally, *D. melanogaster* pigmentation is developmentally plastic, and flies reared at cold temperatures show an increase in melanization (David et al., 1990). Therefore, several lines of evidence suggest that *D. melanogaster* melanization may directly support thermoregulation, similar to various insect and reptile species (Clusella-Trullas et al., 2007). We recently demonstrated that *D. melanogaster* pigmentation evolves rapidly and cyclically across seasons in temperate populations, in which individuals evolve to become darker over winter and lighter in the summer (Berardi et al., 2025). Together, these data suggest temperature drives patterns of pigmentation evolution over space and time.

While temperature and pigmentation have a clear mechanistic link, other aspects of the environment may shape patterns of *D. melanogaster* melanization. Prior work on seasonal pigmentation evolution found an inconsistency: flies remained light late into the fall, despite decreases in daily temperatures (Berardi et al., 2025). This suggests that temperature may not be the sole agent of selection driving melanization patterns, and several additional hypotheses have been put forward to explain the fitness relevance of *D. melanogaster* pigmentation. This includes the proposal that increased melanization provides UV tolerance (Bastide et al., 2014), similar to humans (Jablonski & Chaplin, 2010). Alternatively, melanization may yield structural benefits associated with cuticular strength (Kronforst et al., 2012), and evidence suggests that increased melanization is correlated with improved desiccation tolerance (Rajpurohit et al., 2008b) and immune responses (Nakhleh et al., 2017; Subasi et al., 2024). The canonical *D. melanogaster* pigmentation genes are also highly pleiotropic (Wittkopp & Beldade, 2009); thus, the selective pressures driving change in correlated traits may indirectly shift pigmentation patterns (Rajpurohit et al., 2016). Finally, prior investigation has demonstrated that insect pigmentation can be metabolically costly to produce in resource-limited environments (Ethier et al., 2015; Roff & Fairbairn, 2013), so variation in resource availability and competition may modulate melanization in *Drosophila*. Thus, while temperature remains a likely driver of pigmentation patterns, we can hypothesize that additional agents of selection may directly, or indirectly, impact the seasonal evolution of this complex trait.

A powerful way to test these hypotheses is to manipulate putative environmental drivers of *D. melanogaster* melanization and directly measure evolutionary trajectories of pigmentation. Our recent finding that pigmentation evolves seasonally (Berardi et al., 2025) presented us with the novel opportunity to directly associate rapid evolutionary change in pigmentation with specific shifts in environmental conditions. Thus, to define the selective landscape of *D. melanogaster* pigmentation, we experimentally evolved replicated fly populations across seasons in field mesocosms and manipulated several abiotic and biotic conditions. Our experimental system enabled us to use parallelism across independent replicates to identify adaptive divergence between treatments (Schluter & Nagel, 1995), and to eliminate effects of cryptic structure and/or migration in affecting rapid evolution of pigmentation phenotype. We first tested the hypothesis that temperature is the primary driver of *D. melanogaster* melanization by artificially increasing the temperature in a subset of field mesocosms and assessing whether populations adapted to be less melanized, as predicted by thermal melanism. We then manipulated four additional abiotic and biotic factors that have previously been associated with patterns of seasonal evolution in *Drosophila*: intraspecific competition (Bitter et al., 2024), interspecific competition (Grainger et al., 2019), diet (Beltz et al., 2024), and the gut microbiome (Rudman et al., 2019). Our suite of environmental manipulations allowed us to test whether shifts in melanization appear to be driven solely by temperature to produce a clear fitness benefit (i.e., thermoregulation), or alternatively, if numerous seasonally cycling conditions influence evolutionary patterns. We would predict the latter case to be true if pigmentation is under frequent indirect selection due to correlations with other fitness-related traits that evolve seasonally, yielding a multidimensional selective landscape.

## Materials and Methods

### Experimental orchard system

In order to directly test hypotheses regarding the selective landscape of *Drosophila melanogaster* pigmentation, we designed a series of experiments to examine the effects of targeted environmental manipulations on shifts in melanization. We experimentally evolved replicated *D. melanogaster* populations in field mesocosms (8m^3^ mesh cages) located in Philadelphia, PA, according to the methodology described by: Rajpurohit et al. (2017); Rajpurohit et al. (2018); Rudman et al. (2019); Grainger et al. (2021); Rudman et al. (2022); Bitter et al. (2024); Beltz et al. (2024); Berardi et al. (2025); Karageorgi et al. (2025). The pigmentation data detailed in this paper were compiled across 4 years of experimentation. In each experiment year, we founded field mesocosms with a replicated, outbred *D. melanogaster* starting population. This enabled us to measure parallel shifts in pigmentation from early summer to late fall and determine whether manipulated populations consistently exhibited a differential response relative to controls. By identifying parallel shifts in melanization across numerous replicate populations, and by using our mesocosm system to eliminate confounding effects of migration and demography, we are able to conclude that any significant shifts we observe are most likely adaptive.

Population density was reduced in order to lessen intraspecific competition in two years of experimentation: 2017 and 2022. In 2017, 150 isofemale lines that were originally collected from wild orchard populations in southeastern Pennsylvania were recombined to create an outbred baseline population (Rudman et al., *unpublished data*). We founded our experimental orchard cages with this starting population on June 15. Eight replicate cages were subjected to control conditions (no manipulation, but seasonally evolving), and in an additional eight cages the population density was reduced by removing ∼15% of embryos on the food supply/oviposition substrate every 2d. Flies were then collected from each cage across four timepoints (August 8, September 22, October 18, and November 12) by aspiration and brought into the laboratory for two generations of common garden rearing (25°C, 12L:12D). F3 females were preserved 3-5 days post-eclosion in 75% EtOH at −20°C for later pigmentation scoring; notably, samples of the baseline population for this experiment were lost and not scored for pigmentation.

We separately manipulated intraspecific competition and temperature in 2022. 76 inbred lines originally collected from Media, PA, were recombined across four generations to construct an outbred baseline population (as described in Bitter et al., 2024); this population was then used to found 27 replicate cages (each with 500 males and 500 females from a single 24h, density-controlled cohort) on July 6. We reduced the population density to lessen intraspecific competition in 9 cages, we warmed the temperature in 9 cages, and the remaining 9 cages were unmanipulated controls. Across treatments, populations were provided a fresh loaf pan of standard cornmeal-molasses food every 2 days. To manipulate population density, we reduced the number of eggs laid on the food substrate every 2d to 1,000, which represented low density development conditions based on the ratio of food volume to eggs (Shabalina et al., 1997). When unmanipulated, the egg count in control populations routinely exceeded 10,000 eggs per feeding (every 2d). To manipulate temperature, we artificially warmed a subset of mesocosms by wrapping them in polyethylene greenhouse material to generate an open-top chamber effect (Welshofer et al., 2017). This passively increased the mean daily high temperature in the warming cages by 0.99°C relative to the control treatment during the summer phase, and by 0.73°C during the fall phase (Table S1). We sampled flies from each cage on September 7 and November 8, and we preserved females for pigmentation scoring following two generations of laboratory, common garden treatment as described above.

In 2019, we manipulated interspecific competition between *D. melanogaster* and *Zaprionus indianus* as described in Grainger et al. (2021). This experiment was initiated on July 9, 2019, using an outbred founder population that was established by recombining 150 isofemale lines that were initially collected from wild orchard populations in southeastern Pennsylvania. Here, we focused on the subset of 14 seasonally-evolving cages that either experienced interspecific competition (7 cages) or control conditions (7 cages). During the summer phase of the experiment, *Z. indianus* individuals were introduced to the treatment cages to impose interspecific competition. All flies were removed from the cages the end of the summer, and a subset of 5,000 *D. melanogaster* individuals were reintroduced to each cage to assess the impact of competitive history on subsequent adaptive trajectories during the fall phase. Adult females corresponding to the baseline, summer (September 11), and fall (November 8) timepoints were saved for pigmentation measurements following common garden treatment as described above.

The final two manipulations, diet and food supplementation with specific, gut-colonizing microbes, were conducted in 2020 (Beltz et al., 2024). In this experiment, 80 inbred lines initially collected from southeastern Pennsylvania were recombined to generate an outbred founder population (as described in Rudman et al., 2022). On July 15, we seeded orchard cages with replicates of our founder population to investigate the effects of manipulated diet and microbiome. To test the effects of food quality, we fed flies an apple-based diet containing less protein, fat, and carbohydrate content relative to the standard cornmeal-molasses fly food, as described in Beltz et al., (2024). We then investigated the effects of manipulating the gut microbiome by consistently feeding flies a diet of apple food supplemented with the addition of one of two *D. melanogaster* resident microbes: *Acetobacter thailandicus* and *Lactobacillus brevis* (Beltz et al., 2024; Pais et al., 2018). Treatments were randomly assigned to orchard cages: control food (N=6 cages), apple food (N=6 cages), apple food supplemented with *A. thailandicus* (N=6 cages), and apple food supplemented with *L. brevis* (N=6 cages). Cages were sampled across 5 timepoints in addition to the baseline for the diet treatment (August 13, September 7, September 30, November 9, and November 24), and across 3 timepoints for the microbial addition treatments (September 7, November 9, and November 24). Cage populations were then subjected to two generations of common garden treatment under standardized conditions (25°C, 12L:12D) on the same culture medium they were fed in the field (i.e., control cages were reared on standard food, while the diet and microbe treatments were reared on apple food). Because the microbes were added to apple food, the control for the microbial experiments is the apple treatment populations.

### Scoring abdominal pigmentation

All measurements of abdominal pigmentation were made using a rubric that assigns a score of 1-10 to each abdominal tergite based on the percent melanization (David et al., 1990), where a score of 10 corresponds to 100% melanization. The individual scores for each tergite are then summed to calculate an overall “pigmentation score” for the fly. Preserved females were scored during the same timeframe and blind of sample identity to avoid bias, and the first author quality checked all measurements. We only scored female abdominal pigmentation, as it is more highly variable than male abdominal pigmentation, and all adult females were aged for 3-5 days following eclosure to standardize their development before measuring pigmentation. For all experiment years besides 2017, only the three most distal tergites (tergites 5-7) were scored (David et al., 1990). In 2017 (manipulation of intraspecific competition), all 7 abdominal tergites were scored as in Rajpurohit et al., (2016).

We determined whether each treatment significantly impacted pigmentation patterns by running a linear mixed effects model in R (RStudio v.2024.04.1+748) using the function ‘lmer’ in the package ‘lme4’ (v.1.1.35.5), and significance was assessed with an ANOVA (‘lmerTest’ v.3.1.3). For each experiment, timepoint, treatment, and their interaction were modeled as fixed effects, and cage was included as a random effect per the formula “lmer(Pigmentation Score ∼ Timepoint * Treatment + (1|Cage)”. For each model, we then ran planned comparisons to test (i) differences between treatment and control pigmentation at the end of the summer and fall phases and (ii) significant shifts in pigmentation from the summer to fall for each treatment. Here, we used the function ‘contrast’ from the package ‘emmeans’ (v.1.10.2) to run *t-*tests across each planned comparison; we have reported both raw and Holm corrected *p-*values (Tables S2-S8). The data for each experiment was plotted using ‘ggplot2’ (v.3.5.1), and the R packages ‘tidyverse’ (v.2.0.0), ‘readxl’ (v.1.4.3), and ‘reshape2’ (v.1.4.4) were used in data analysis. Figures were made using Keynote (v.8.1). Raw data and scripts associated with the analyses for this project can be accessed at: https://github.com/skylerberardi/Papers/tree/main/Berardi_pigmentation.drivers

## Results

### Temperature drives differential pigmentation patterns during the summer

We first tested whether temperature is a significant driver of seasonal pigmentation patterns in *D. melanogaster*, and we found that increasing the temperature altered the adaptive trajectory of melanization over time relative to the control (Fig. 1; Table S2A; *F* = 7.32, *p* = .007). The warming treatment drove populations to exhibit significantly lighter pigmentation following the summer phase of seasonal evolution (Table S2B; *t* = 3.20, *p* = .007), and this result was concordant with the prediction that reduced melanization would be favored under warmer temperatures to prevent overheating under a thermoregulatory model (Freoa et al., 2023). However, pigmentation patterns were not aligned with the thermal melanism hypothesis during the fall phase of the experiment. As daily temperatures dropped, we expected both the warming and control populations to evolve darker pigmentation, and the control populations to be the most melanized. Instead, we observed that the control populations evolved to be lighter in the fall, as seen in previous work (Berardi et al., 2025), and melanization did not significantly differ between treatments (Fig. 1; Table S2B). Thus, our findings support the hypothesis that temperature is a key driver of pigmentation patterns during the summer phase, but not during the fall phase. This suggests that the seasonal selective landscape for pigmentation is temporally variable and complex, and we aimed to uncover whether other key components of the environment influence adaptive pigmentation patterns.

**Figure 1.**
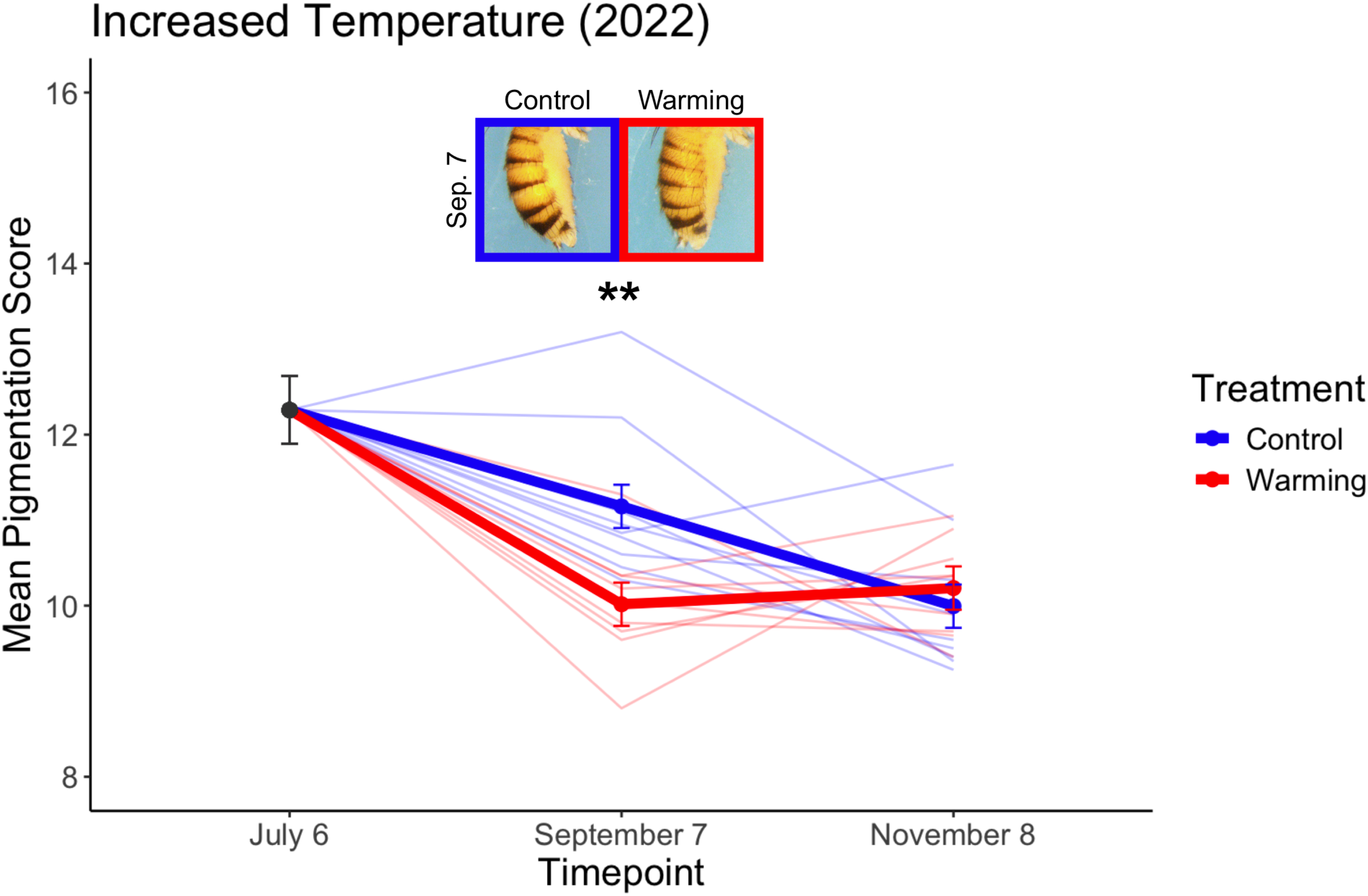
Increasing the temperature reduces melanization during the summer phase. (A) Populations subjected to our ‘warming’ treatment exhibited significantly lighter pigmentation relative to the control populations following the summer phase, but the populations did not differ in melanization following the fall phase of evolution. In all plots, mean pigmentation score across each treatment (+/− SE) are plotted alongside individual cage scores (thin lines). Representative images depicting the mean pigmentation score for the control and warming treatments are shown for the summer phase (\*\**p* < .01).

### Intraspecific competition influences the adaptive trajectory of pigmentation

Fluctuations in intraspecific competition associated with seasonal boom-bust dynamics (Bitter et al., 2024; Gleason et al., 2019), as well as seasonal shifts in the extent of interspecific competition (Grainger at al., 2021), have been associated with patterns of rapid evolution in *D. melanogaster*. Therefore, we first aimed to test whether modulating levels of intra- and interspecific competition drive differential pigmentation patterns across seasons. To manipulate the magnitude of intraspecific competition, we reduced *D. melanogaster* population density in two separate years of experimentation (2017 and 2022) and compared evolutionary trajectories of pigmentation to control cages where density was unmanipulated. In both years, reducing the population density yielded the same effect on pigmentation patterns over time: populations with reduced intraspecific competition evolved to be significantly darker in the fall relative to control populations (light lines in Fig. 2A,B). We measured pigmentation patterns at four timepoints in 2017, and we found that melanization varied significantly by treatment (Fig. 2A; Table S3A; *F* = 49.38, *p* < .001), and the effects of treatment varied with time (*F =* 94.30, *p* < .001). Contrasts showed a significant difference between the control and density treatments at the ends of both the summer (*t* = −2.82, *p* = .007) and the fall phases (*t =* −15.20, *p* < .001; Table S3B). In 2022, we similarly saw that the reduced density treatment altered the evolutionary trajectory of pigmentation patterns relative to control (Fig. 2B; Table S4A; *F* = 17.02, *p* < .001), and reduced density treatment populations were significantly darker than control populations after the fall phase of evolution (Table S4B; *t =* −3.53, *p* = .004).

**Figure 2.**
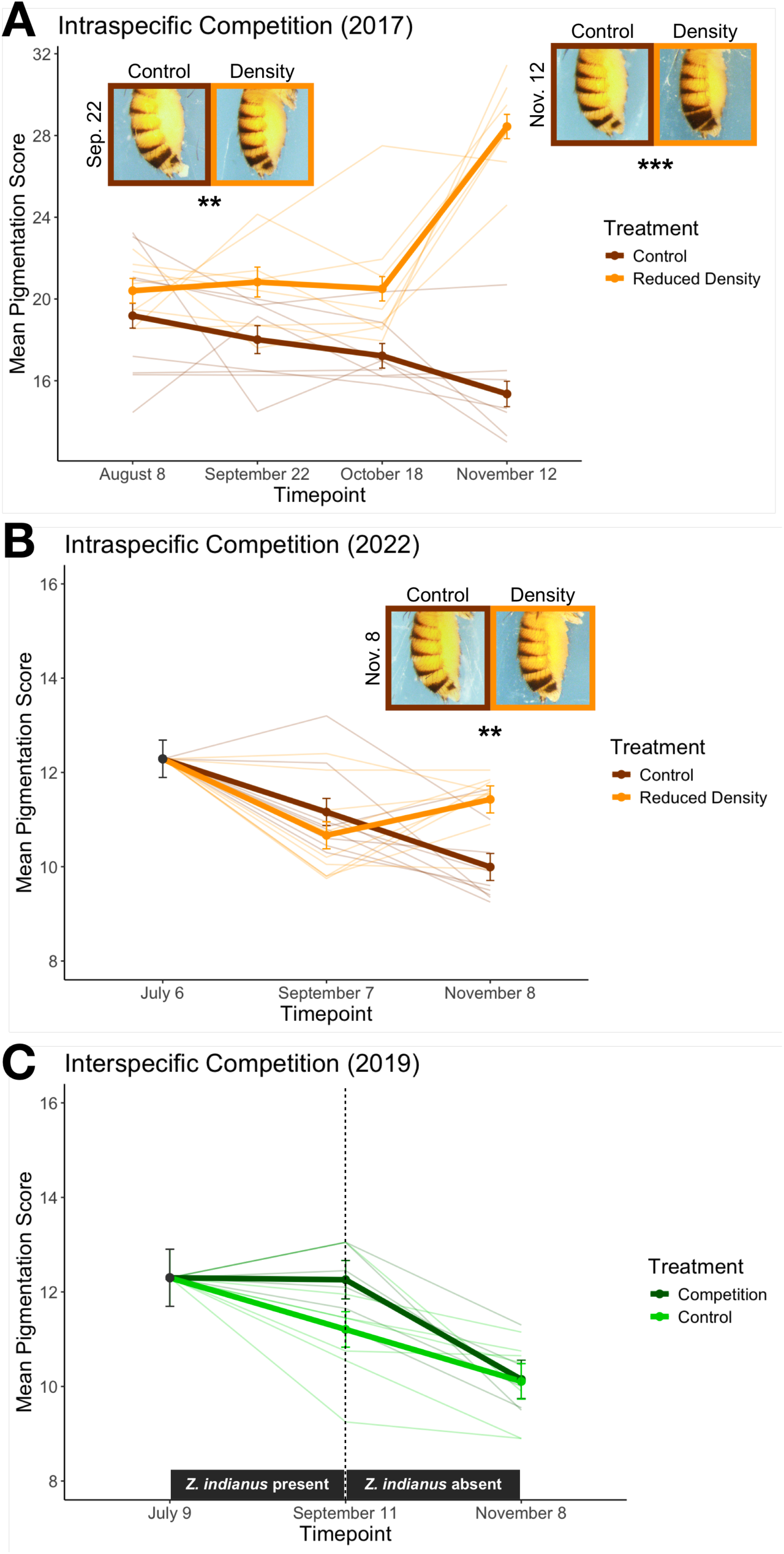
Manipulations of intraspecific competition drive differential pigmentation patterns across seasons. (A,B) Reducing *D. melanogaster* population density in two separate years of experimentation drove darker pigmentation relative to the control following the fall phase. Notably, all seven abdominal tergites were scored for the reduced density treatment in 2017 (panel B; see Methods), yielding a different y-axis scale. (C) Introducing *Z. indianus* as an interspecific competitor during the summer phase was associated with a nonsignificant increase in melanization. *Z. indianus* individuals were then removed from each competition treatment cage during the fall phase. In all plots, mean pigmentation score across each treatment (+/− SE) are plotted alongside individual cage scores (thin lines). Representative images depicting the population mean are shown at the end of the summer and/or fall phase if differences in melanization between treatments are significant (\*\*\**p* < .001, \*\**p* < .01).

In addition to manipulating intraspecific competition, we also tested for effects of interspecific competition on pigmentation evolution by introducing a competitor fly, *Zaprionus indianus*, to a subset of field mesocosms (Grainger et al., 2021). However, our control and interspecific competition populations did not exhibit significantly different pigmentation patterns (Fig. 2C; Table S5A). We found that introducing *Z. indianus* during the summer phase of the experiment led to a nonsignificant increase in melanization at our September timepoint relative to the control (Fig. 2C; Table S5B; *t* = −1.90, *p* = .136). *Z. indianus* were absent from the competition treatment mesocosms during the fall phase, and we observed that both the control and treatment populations evolved to be significantly lighter in the fall relative to the summer but did not differ in pigmentation (Table S5B).

### Manipulations of diet and the gut microbiome yield differential shifts in melanization

Previous work has also demonstrated that diet (Beltz et al., 2024) and the gut microbiome (Rudman et al., 2019) represent key drivers of seasonal adaptation in *D. melanogaster*, and we examined whether these factors influence the evolution of pigmentation. We first found that feeding flies a lower quality, apple-based diet (Beltz et al., 2024) altered the seasonal trajectory of pigmentation adaptation relative to the higher quality control diet (Fig. 3A; Table S6A; *F* = 4.14, *p* = .003). Populations evolving on the apple diet exhibited darker pigmentation relative to the control diet at the end of the summer (*t* = −2.49, *p* = .028) and fall phases (*t* = −2.31, *p* = .028; Table S6B), but patterns of melanization fluctuated between treatments at intermediate timepoints (Fig. 3A). We then altered the gut microbiome of *D. melanogaster* populations by supplementing the apple diet with one of two resident microbes: *A. thailandicus* and *L. brevis* (Fig. 3B). We found that treating populations with either *A. thailandicus* (Table S7A; *F* = 4.73, *p* = .009) or *L. brevis* (Table S8A; *F* = 8.37, *p* < .001) altered the adaptive trajectory of melanization over time relative to control. Interestingly, the effects on pigmentation patterns at specific timepoints differed for each microbe (Fig. 3B; Table S7B; Table S8B).

**Figure 3.**
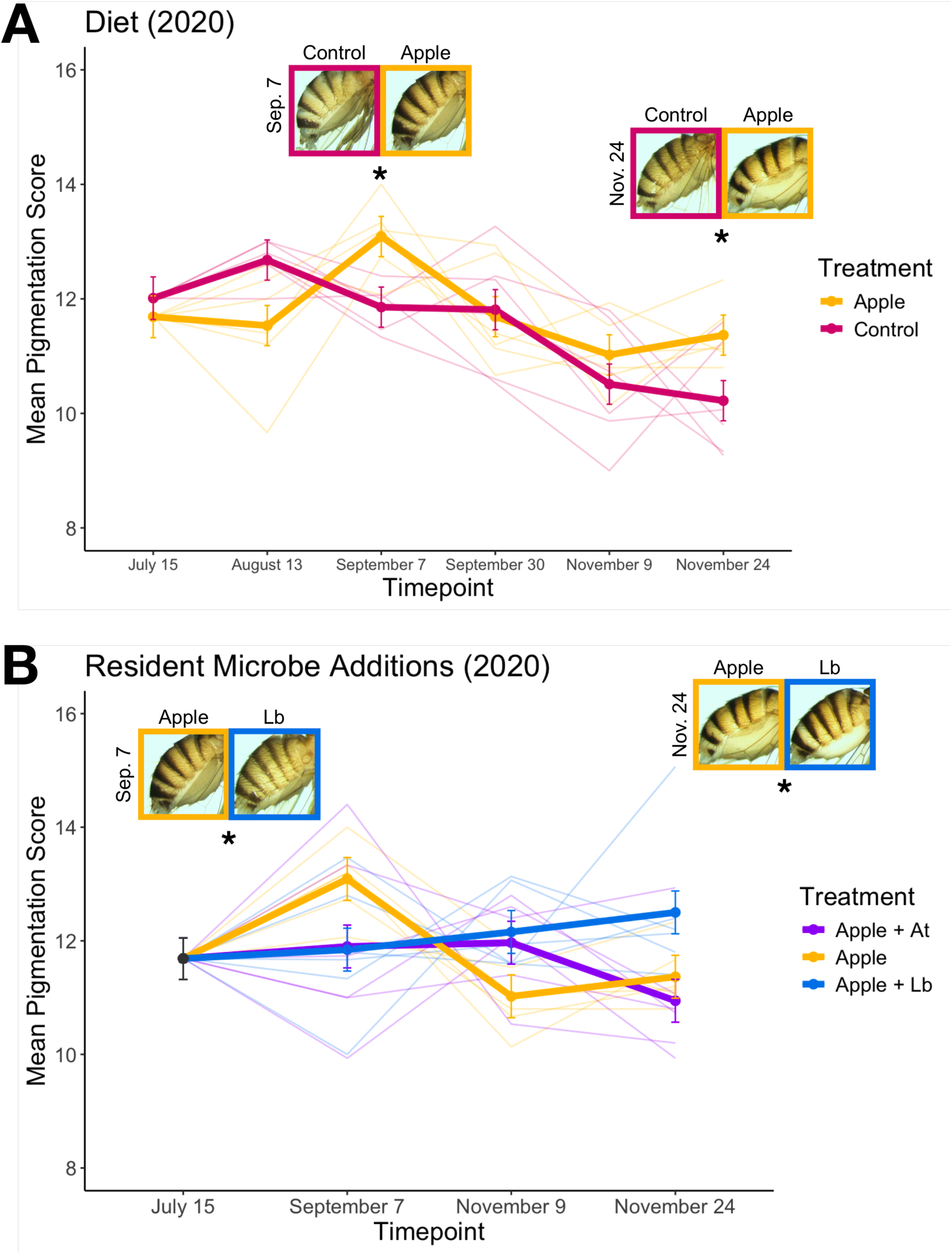
Distinct pigmentation patterns arise following manipulations of diet and the gut microbiome. (A) Populations subjected to our apple diet treatment exhibited shifts in melanization over time that differed significantly to shifts in the control populations. (B) Both resident microbe additions (*A. thailandicus* and *L. brevis*) produced melanization patterns that varied across seasons relative to control populations. Here, the control populations are the apple-based food treatment (shown in A), as apple food was supplemented with each microbe in this experiment. The *A. thailandicus* treatment is denoted as “Apple + At”, and the *L. brevis* treatment is denoted as “Apple + Lb”. In all plots, the mean pigmentation score across each treatment (+/− SE) is shown alongside individual cage scores (thin lines). Representative images depicting the population mean for each treatment are shown at the end of the summer and fall (\**p* < .05).

## Discussion

We utilized a field-based, experimental evolution approach to examine how multiple environmental agents of selection influence the adaptive dynamics of pigmentation over ecological timescales (Hairston et al., 2005). We first observed that our unmanipulated control populations consistently evolved to be less melanized over time, showing a parallel response across all four years of experimentation (Tables S2B-S8B) despite inter-year variation in the composition of our outbred founding population (see Methods). This finding was concordant with our expectations based on previous work, which demonstrated that temperate *D. melanogaster* populations repeatedly evolve lighter pigmentation following the summer growing season (Berardi et al., 2025). We then explored the complexity of the seasonal selective landscape for pigmentation to identify the environmental agents that drive parallel evolutionary patterns indicative of adaptation. Under the simplest scenario, we anticipated that patterns of melanization would respond to manipulations of temperature, in line with the thermal melanism hypothesis (Clusella-Trullas et al., 2007). However, while we confirmed that increasing the temperature drove lighter pigmentation during the summer, as would be predicted if lessening melanization minimizes overheating, we did not observe a shift to darker pigmentation to promote heating when conditions became colder during the fall. Overall, we found that manipulating several other key environmental parameters drove differential pigmentation patterns over seasons. Thus, our findings illustrate that the selective landscape of *D. melanogaster* pigmentation is multifaceted, and temperature gradients alone may not be sufficient to explain melanization patterns in wild populations.

Both the parallelism in the evolutionary response across replicates, and our ability to control for confounding effects of migration and demography with our field mesocosm system, enable us to conclude that the changes in pigmentation patterns we observed are most likely adaptive. However, whether pigmentation is under direct or indirect selection under each set of conditions remains an open question that should be addressed through future lines of inquiry. Our temperature manipulation provided some evidence that pigmentation may be under direct selection to produce a thermoregulatory benefit during the summer. We observed that warming treatment populations evolved to be lighter than control populations following the summer months, and this is concordant with our expectations if lighter pigmentation produces a cooler body temperature in the field (Freoa et al., 2023). This finding also aligns with a breadth of previous work showing that lighter pigmentation covaries with warmer temperatures over latitudinal (Telonis-Scott et al., 2011) and altitudinal (Bastide et al., 2014), and seasonal gradients (Berardi et al., 2025) in wild populations. However, we did not measure the body temperature of flies in our warming and control treatments, and we also did not observe increased melanization as temperatures decreased later in the fall, as would be predicted by the thermal melanism hypothesis. Unlike other systems where variation in pigmentation can be experimentally created to directly test fitness consequences (e.g., Kingsolver, 1987), laboratory validation of the thermal melanism hypothesis remains challenging in *Drosophila.* Therefore, we cannot presently conclude whether *D. melanogaster* melanization directly supports thermoregulation.

Notably, we found that pigmentation patterns evolved in response to components of the seasonal environment that were previously not connected to melanization in *D. melanogaster*: intraspecific competition, diet, and the gut microbiome. This suggests that evolution of pigmentation is either a component of adaptation to a wide range of stressors or is indirectly driven by correlations with other traits under seasonally varying selection. The loci that regulate insect pigmentation are notoriously pleiotropic (Wittkopp & Beldade, 2009), and *D. melanogaster* pigmentation genes have been implicated in affecting mating behavior (Drapeau et al., 2003; Wilson et al., 1976), morphology (Godt et al., 1993; Kopp et al., 2000), cuticle hardening and immunity (Hodgetts & O’Keefe, 2006; Jacobs, 1985), circadian activity (Shaw et al., 2000), vision (True et al., 2005), and cuticular hydrocarbon composition (Massey et al., 2019). We know that numerous fitness-related phenotypes evolve seasonally in *D. melanogaster* (Behrman et al., 2015; Erickson et al., 2020; Rudman et al., 2022; Schmidt & Conde, 2006), including traits with pleiotropic associations to pigmentation such as innate immunity (Behrman et al., 2018), cuticular hydrocarbon content (Rajpurohit et al., 2017), and desiccation tolerance (Rajpurohit et al., 2018). Therefore, pigmentation may be subject to frequent indirect selection due to genetic correlations, and this provides a reasonable explanation for its responsiveness to a breadth of ecological manipulations.

Interestingly, both intraspecific competition manipulations yielded pigmentation shifts as predicted under the thermal melanism model: reducing the population density drove lighter pigmentation in the summer, and darker pigmentation in the fall. There is evidence that melanin is metabolically costly to produce across both insects (Ethier et al., 2015; Roff & Fairbairn, 2013; Stoehr 2006) and vertebrates (Britton & Davidowitz, 2023; Stoehr 2006), which provides a possible explanation for this pattern. If melanin synthesis is costly under stressful conditions, alleviating intraspecific competition for resources should yield a higher capacity to produce melanin, as observed. Conversely, melanization may be constrained by increased competition in the control populations, imposing a metabolic trade-off that limits the synthesis of pigment. Therefore, it is possible that lessening competition reduced fitness costs of melanin production, leading to flies evolving darker pigmentation and improving thermoregulation in the fall. Importantly, however, the results of our diet manipulation were not aligned with the hypothesis that melanin is costly in resource limited environments. The apple diet represented a reduction in nutritional quality relative to the control diet (Beltz et al., 2024); thus, if melanin is costly to produce, we would have anticipated flies reared on apple food to consistently be lighter than flies reared on control food across all seasonal timepoints, which was not the case. We also failed to see any significant change in pigmentation patterns in our interspecific competition treatment relative to the control, despite the resource environment varying between treatments. Therefore, further investigation into metabolic trade-offs of melanization in *Drosophila* is required to fully interpret our findings.

Altogether, we demonstrate that the adaptive dynamics of *D. melanogaster* pigmentation are far more nuanced than previously appreciated, which provides exciting opportunities for future exploration. It is clear that temperature is not the sole driver of pigmentation in *D. melanogaster*, so revisiting melanization patterns in the context of life history theory and energetic trade-offs represent important steps for yielding insight into the determinants of phenotype. Pigmentation patterns were highly responsive to a wide range of environmental manipulations, and defining whether these shifts are largely driven by direct fitness benefits or indirect selection remains a key area of investigation. Future work can also utilize *D. melanogaster* pigmentation as a tool to explore how differential shifts at the genetic level are correlated with patterns of melanization across varying environments. Finally, examining the agents of selection acting on other complex traits will enable us to understand whether the selective landscape for pigmentation is uniquely multifaceted, or if it represents the standard for how patterns of complex trait adaptation are shaped in dynamic environments.

## Author Contributions

S.B. and P.S. conceptualized and designed the overall study. S.M.R. and P.S. designed and conducted the 2017 orchard experiment. T.N.G., J.M.L., S.M.R., and P.S. designed and conducted the 2019 orchard experiment. J.K.B. and P.S. designed and conducted the 2020 orchard experiment. S.B. and P.S. designed and conducted the 2022 orchard experiment. H.O. contributed to field work for the 2019, 2020, and 2022 experiments, and J.K.B. contributed to field work for the 2022 experiment. S.B. led measurement and analysis of pigmentation data. S.B. and P.S. wrote the manuscript, with feedback provided by all authors.

## Acknowledgements

We thank current and past members of the P.S. research group for their feedback on this manuscript.

## Data and Code Availability

Raw data and scripts associated with this project have been uploaded to the following Github repository: https://github.com/skylerberardi/Papers/tree/main/Berardi_pigmentation.drivers

## Supplementary Materials

**Table S1.**
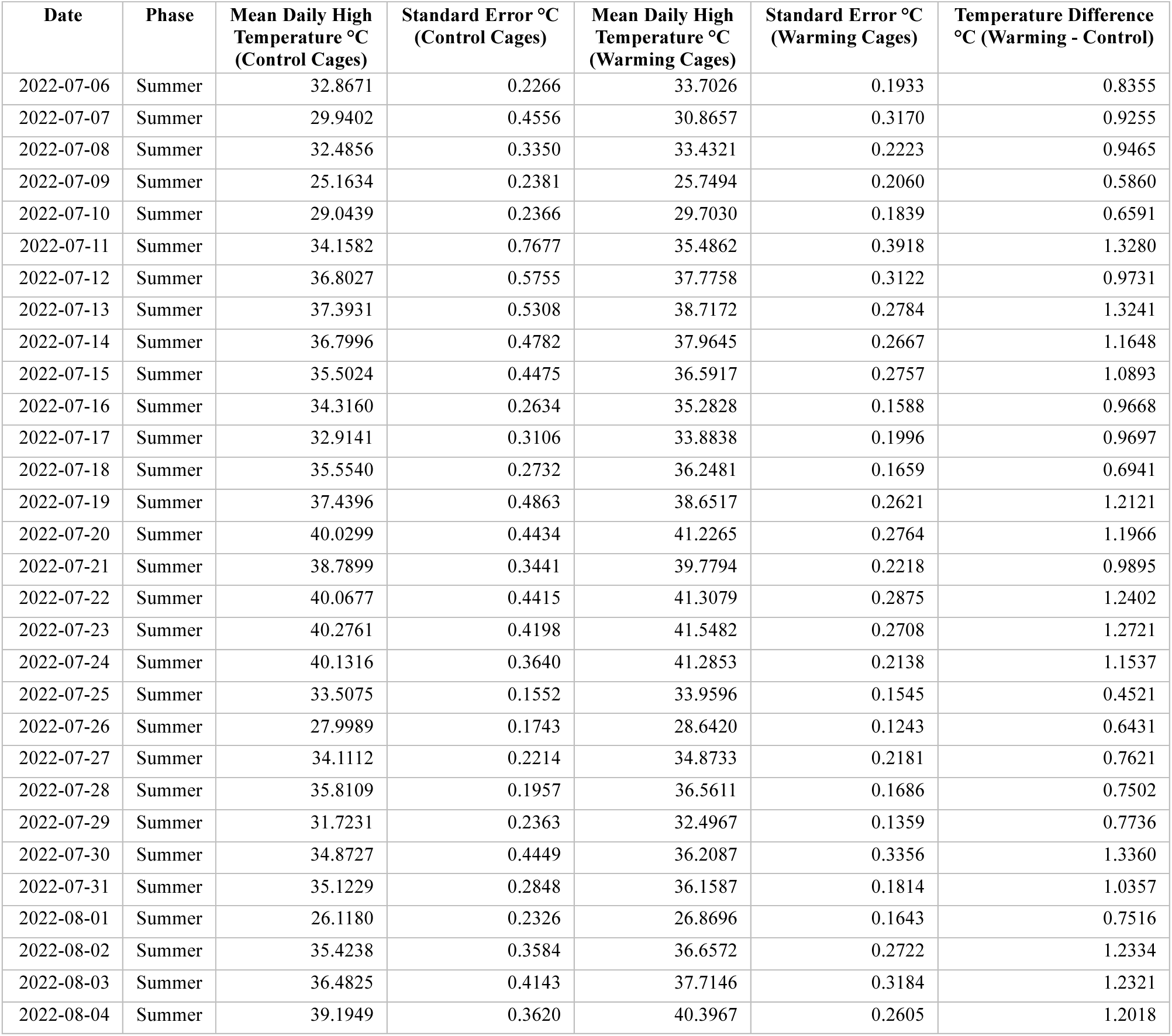
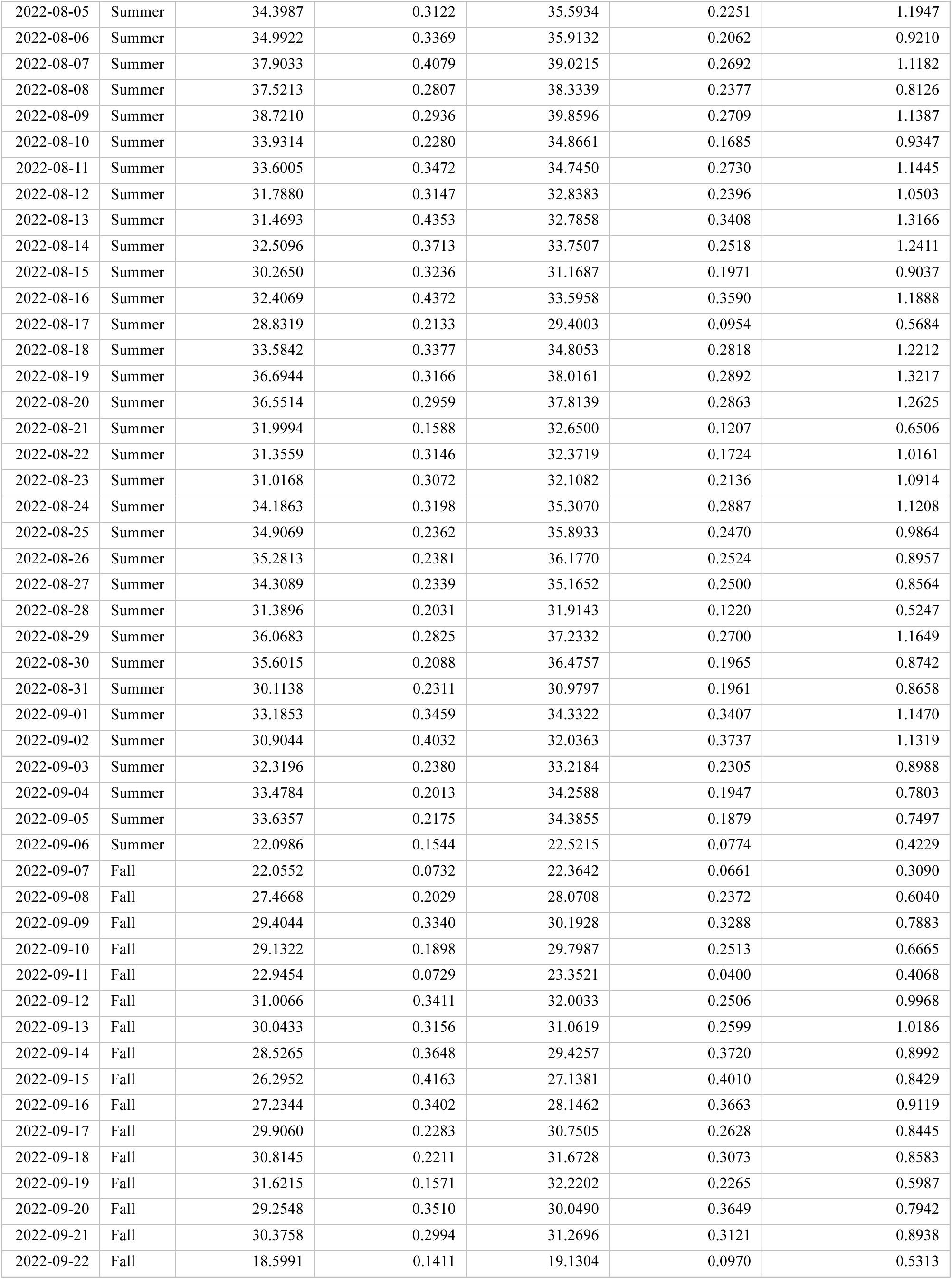
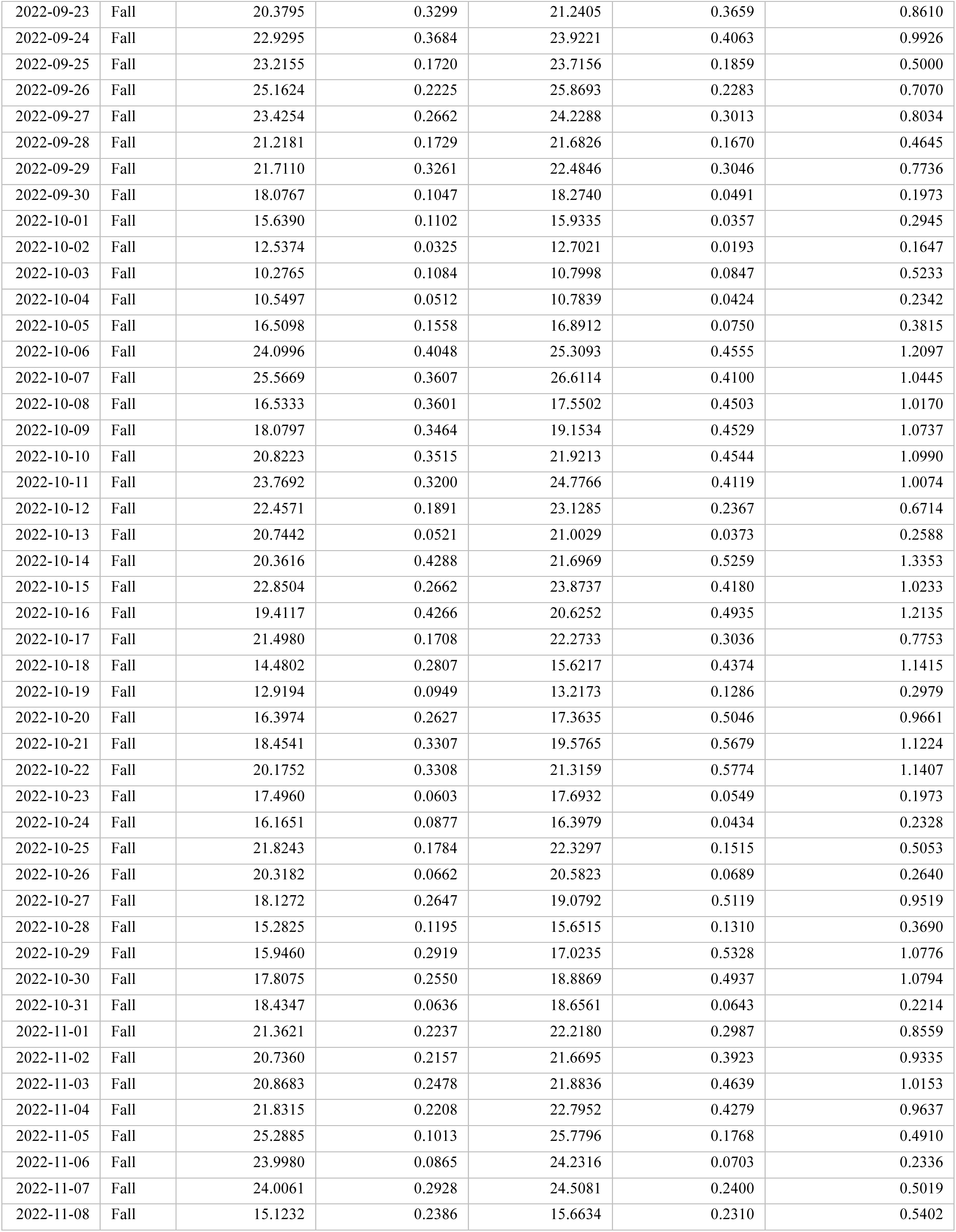
Mean daily high temperatures across control and warming cages. Temperature measurements were taken with data loggers (HOBO Pendant MX2202) throughout the 2022 experiment at 10-minute intervals, and the daily high temperature for each cage was calculated by averaging temperatures recorded from 2:00 PM – 4:00 PM. Two outlier control cages (“E2” and “E3”) were removed from the analysis; the peach trees in these cages were replaced at the beginning of the season and provided less shade, and the cages exhibited elevated temperatures relative to the other control cages. The mean daily high temperature was calculated at each date across the control cages and the warming cages, and we then determined the temperature difference at each date between the warming and control treatments. We found that across the entire season, mean daily high temperatures in the warming treatment were 0.86 +/− 0.03°C greater than the control. This difference was 0.99 +/− 0.03°C across the summer phase, and 0.73 +/− 0.04°C across the fall phase (mean +/− standard error).

| Date | Phase | Mean Daily High Temperature $^{\circ}\text{C}$ (Control Cages) | Standard Error $^{\circ}\text{C}$ (Control Cages) | Mean Daily High Temperature $^{\circ}\text{C}$ (Warming Cages) | Standard Error $^{\circ}\text{C}$ (Warming Cages) | Temperature Difference $^{\circ}\text{C}$ (Warming - Control) |
| --- | --- | --- | --- | --- | --- | --- |
| 2022-07-06 | Summer | 32.8671 | 0.2266 | 33.7026 | 0.1933 | 0.8355 |
| 2022-07-07 | Summer | 29.9402 | 0.4556 | 30.8657 | 0.3170 | 0.9255 |
| 2022-07-08 | Summer | 32.4856 | 0.3350 | 33.4321 | 0.2223 | 0.9465 |
| 2022-07-09 | Summer | 25.1634 | 0.2381 | 25.7494 | 0.2060 | 0.5860 |
| 2022-07-10 | Summer | 29.0439 | 0.2366 | 29.7030 | 0.1839 | 0.6591 |
| 2022-07-11 | Summer | 34.1582 | 0.7677 | 35.4862 | 0.3918 | 1.3280 |
| 2022-07-12 | Summer | 36.8027 | 0.5755 | 37.7758 | 0.3122 | 0.9731 |
| 2022-07-13 | Summer | 37.3931 | 0.5308 | 38.7172 | 0.2784 | 1.3241 |
| 2022-07-14 | Summer | 36.7996 | 0.4782 | 37.9645 | 0.2667 | 1.1648 |
| 2022-07-15 | Summer | 35.5024 | 0.4475 | 36.5917 | 0.2757 | 1.0893 |
| 2022-07-16 | Summer | 34.3160 | 0.2634 | 35.2828 | 0.1588 | 0.9668 |
| 2022-07-17 | Summer | 32.9141 | 0.3106 | 33.8838 | 0.1996 | 0.9697 |
| 2022-07-18 | Summer | 35.5540 | 0.2732 | 36.2481 | 0.1659 | 0.6941 |
| 2022-07-19 | Summer | 37.4396 | 0.4863 | 38.6517 | 0.2621 | 1.2121 |
| 2022-07-20 | Summer | 40.0299 | 0.4434 | 41.2265 | 0.2764 | 1.1966 |
| 2022-07-21 | Summer | 38.7899 | 0.3441 | 39.7794 | 0.2218 | 0.9895 |
| 2022-07-22 | Summer | 40.0677 | 0.4415 | 41.3079 | 0.2875 | 1.2402 |
| 2022-07-23 | Summer | 40.2761 | 0.4198 | 41.5482 | 0.2708 | 1.2721 |
| 2022-07-24 | Summer | 40.1316 | 0.3640 | 41.2853 | 0.2138 | 1.1537 |
| 2022-07-25 | Summer | 33.5075 | 0.1552 | 33.9596 | 0.1545 | 0.4521 |
| 2022-07-26 | Summer | 27.9989 | 0.1743 | 28.6420 | 0.1243 | 0.6431 |
| 2022-07-27 | Summer | 34.1112 | 0.2214 | 34.8733 | 0.2181 | 0.7621 |
| 2022-07-28 | Summer | 35.8109 | 0.1957 | 36.5611 | 0.1686 | 0.7502 |
| 2022-07-29 | Summer | 31.7231 | 0.2363 | 32.4967 | 0.1359 | 0.7736 |
| 2022-07-30 | Summer | 34.8727 | 0.4449 | 36.2087 | 0.3356 | 1.3360 |
| 2022-07-31 | Summer | 35.1229 | 0.2848 | 36.1587 | 0.1814 | 1.0357 |
| 2022-08-01 | Summer | 26.1180 | 0.2326 | 26.8696 | 0.1643 | 0.7516 |
| 2022-08-02 | Summer | 35.4238 | 0.3584 | 36.6572 | 0.2722 | 1.2334 |
| 2022-08-03 | Summer | 36.4825 | 0.4143 | 37.7146 | 0.3184 | 1.2321 |
| 2022-08-04 | Summer | 39.1949 | 0.3620 | 40.3967 | 0.2605 | 1.2018 |
| 2022-08-05 | Summer | 34.3987 | 0.3122 | 35.5934 | 0.2251 | 1.1947 |
| 2022-08-06 | Summer | 34.9922 | 0.3369 | 35.9132 | 0.2062 | 0.9210 |
| 2022-08-07 | Summer | 37.9033 | 0.4079 | 39.0215 | 0.2692 | 1.1182 |
| 2022-08-08 | Summer | 37.5213 | 0.2807 | 38.3339 | 0.2377 | 0.8126 |
| 2022-08-09 | Summer | 38.7210 | 0.2936 | 39.8596 | 0.2709 | 1.1387 |
| 2022-08-10 | Summer | 33.9314 | 0.2280 | 34.8661 | 0.1685 | 0.9347 |
| 2022-08-11 | Summer | 33.6005 | 0.3472 | 34.7450 | 0.2730 | 1.1445 |
| 2022-08-12 | Summer | 31.7880 | 0.3147 | 32.8383 | 0.2396 | 1.0503 |
| 2022-08-13 | Summer | 31.4693 | 0.4353 | 32.7858 | 0.3408 | 1.3166 |
| 2022-08-14 | Summer | 32.5096 | 0.3713 | 33.7507 | 0.2518 | 1.2411 |
| 2022-08-15 | Summer | 30.2650 | 0.3236 | 31.1687 | 0.1971 | 0.9037 |
| 2022-08-16 | Summer | 32.4069 | 0.4372 | 33.5958 | 0.3590 | 1.1888 |
| 2022-08-17 | Summer | 28.8319 | 0.2133 | 29.4003 | 0.0954 | 0.5684 |
| 2022-08-18 | Summer | 33.5842 | 0.3377 | 34.8053 | 0.2818 | 1.2212 |
| 2022-08-19 | Summer | 36.6944 | 0.3166 | 38.0161 | 0.2892 | 1.3217 |
| 2022-08-20 | Summer | 36.5514 | 0.2959 | 37.8139 | 0.2863 | 1.2625 |
| 2022-08-21 | Summer | 31.9994 | 0.1588 | 32.6500 | 0.1207 | 0.6506 |
| 2022-08-22 | Summer | 31.3559 | 0.3146 | 32.3719 | 0.1724 | 1.0161 |
| 2022-08-23 | Summer | 31.0168 | 0.3072 | 32.1082 | 0.2136 | 1.0914 |
| 2022-08-24 | Summer | 34.1863 | 0.3198 | 35.3070 | 0.2887 | 1.1208 |
| 2022-08-25 | Summer | 34.9069 | 0.2362 | 35.8933 | 0.2470 | 0.9864 |
| 2022-08-26 | Summer | 35.2813 | 0.2381 | 36.1770 | 0.2524 | 0.8957 |
| 2022-08-27 | Summer | 34.3089 | 0.2339 | 35.1652 | 0.2500 | 0.8564 |
| 2022-08-28 | Summer | 31.3896 | 0.2031 | 31.9143 | 0.1220 | 0.5247 |
| 2022-08-29 | Summer | 36.0683 | 0.2825 | 37.2332 | 0.2700 | 1.1649 |
| 2022-08-30 | Summer | 35.6015 | 0.2088 | 36.4757 | 0.1965 | 0.8742 |
| 2022-08-31 | Summer | 30.1138 | 0.2311 | 30.9797 | 0.1961 | 0.8658 |
| 2022-09-01 | Summer | 33.1853 | 0.3459 | 34.3322 | 0.3407 | 1.1470 |
| 2022-09-02 | Summer | 30.9044 | 0.4032 | 32.0363 | 0.3737 | 1.1319 |
| 2022-09-03 | Summer | 32.3196 | 0.2380 | 33.2184 | 0.2305 | 0.8988 |
| 2022-09-04 | Summer | 33.4784 | 0.2013 | 34.2588 | 0.1947 | 0.7803 |
| 2022-09-05 | Summer | 33.6357 | 0.2175 | 34.3855 | 0.1879 | 0.7497 |
| 2022-09-06 | Summer | 22.0986 | 0.1544 | 22.5215 | 0.0774 | 0.4229 |
| 2022-09-07 | Fall | 22.0552 | 0.0732 | 22.3642 | 0.0661 | 0.3090 |
| 2022-09-08 | Fall | 27.4668 | 0.2029 | 28.0708 | 0.2372 | 0.6040 |
| 2022-09-09 | Fall | 29.4044 | 0.3340 | 30.1928 | 0.3288 | 0.7883 |
| 2022-09-10 | Fall | 29.1322 | 0.1898 | 29.7987 | 0.2513 | 0.6665 |
| 2022-09-11 | Fall | 22.9454 | 0.0729 | 23.3521 | 0.0400 | 0.4068 |
| 2022-09-12 | Fall | 31.0066 | 0.3411 | 32.0033 | 0.2506 | 0.9968 |
| 2022-09-13 | Fall | 30.0433 | 0.3156 | 31.0619 | 0.2599 | 1.0186 |
| 2022-09-14 | Fall | 28.5265 | 0.3648 | 29.4257 | 0.3720 | 0.8992 |
| 2022-09-15 | Fall | 26.2952 | 0.4163 | 27.1381 | 0.4010 | 0.8429 |
| 2022-09-16 | Fall | 27.2344 | 0.3402 | 28.1462 | 0.3663 | 0.9119 |
| 2022-09-17 | Fall | 29.9060 | 0.2283 | 30.7505 | 0.2628 | 0.8445 |
| 2022-09-18 | Fall | 30.8145 | 0.2211 | 31.6728 | 0.3073 | 0.8583 |
| 2022-09-19 | Fall | 31.6215 | 0.1571 | 32.2202 | 0.2265 | 0.5987 |
| 2022-09-20 | Fall | 29.2548 | 0.3510 | 30.0490 | 0.3649 | 0.7942 |
| 2022-09-21 | Fall | 30.3758 | 0.2994 | 31.2696 | 0.3121 | 0.8938 |
| 2022-09-22 | Fall | 18.5991 | 0.1411 | 19.1304 | 0.0970 | 0.5313 |
| 2022-09-23 | Fall | 20.3795 | 0.3299 | 21.2405 | 0.3659 | 0.8610 |
| 2022-09-24 | Fall | 22.9295 | 0.3684 | 23.9221 | 0.4063 | 0.9926 |
| 2022-09-25 | Fall | 23.2155 | 0.1720 | 23.7156 | 0.1859 | 0.5000 |
| 2022-09-26 | Fall | 25.1624 | 0.2225 | 25.8693 | 0.2283 | 0.7070 |
| 2022-09-27 | Fall | 23.4254 | 0.2662 | 24.2288 | 0.3013 | 0.8034 |
| 2022-09-28 | Fall | 21.2181 | 0.1729 | 21.6826 | 0.1670 | 0.4645 |
| 2022-09-29 | Fall | 21.7110 | 0.3261 | 22.4846 | 0.3046 | 0.7736 |
| 2022-09-30 | Fall | 18.0767 | 0.1047 | 18.2740 | 0.0491 | 0.1973 |
| 2022-10-01 | Fall | 15.6390 | 0.1102 | 15.9335 | 0.0357 | 0.2945 |
| 2022-10-02 | Fall | 12.5374 | 0.0325 | 12.7021 | 0.0193 | 0.1647 |
| 2022-10-03 | Fall | 10.2765 | 0.1084 | 10.7998 | 0.0847 | 0.5233 |
| 2022-10-04 | Fall | 10.5497 | 0.0512 | 10.7839 | 0.0424 | 0.2342 |
| 2022-10-05 | Fall | 16.5098 | 0.1558 | 16.8912 | 0.0750 | 0.3815 |
| 2022-10-06 | Fall | 24.0996 | 0.4048 | 25.3093 | 0.4555 | 1.2097 |
| 2022-10-07 | Fall | 25.5669 | 0.3607 | 26.6114 | 0.4100 | 1.0445 |
| 2022-10-08 | Fall | 16.5333 | 0.3601 | 17.5502 | 0.4503 | 1.0170 |
| 2022-10-09 | Fall | 18.0797 | 0.3464 | 19.1534 | 0.4529 | 1.0737 |
| 2022-10-10 | Fall | 20.8223 | 0.3515 | 21.9213 | 0.4544 | 1.0990 |
| 2022-10-11 | Fall | 23.7692 | 0.3200 | 24.7766 | 0.4119 | 1.0074 |
| 2022-10-12 | Fall | 22.4571 | 0.1891 | 23.1285 | 0.2367 | 0.6714 |
| 2022-10-13 | Fall | 20.7442 | 0.0521 | 21.0029 | 0.0373 | 0.2588 |
| 2022-10-14 | Fall | 20.3616 | 0.4288 | 21.6969 | 0.5259 | 1.3353 |
| 2022-10-15 | Fall | 22.8504 | 0.2662 | 23.8737 | 0.4180 | 1.0233 |
| 2022-10-16 | Fall | 19.4117 | 0.4266 | 20.6252 | 0.4935 | 1.2135 |
| 2022-10-17 | Fall | 21.4980 | 0.1708 | 22.2733 | 0.3036 | 0.7753 |
| 2022-10-18 | Fall | 14.4802 | 0.2807 | 15.6217 | 0.4374 | 1.1415 |
| 2022-10-19 | Fall | 12.9194 | 0.0949 | 13.2173 | 0.1286 | 0.2979 |
| 2022-10-20 | Fall | 16.3974 | 0.2627 | 17.3635 | 0.5046 | 0.9661 |
| 2022-10-21 | Fall | 18.4541 | 0.3307 | 19.5765 | 0.5679 | 1.1224 |
| 2022-10-22 | Fall | 20.1752 | 0.3308 | 21.3159 | 0.5774 | 1.1407 |
| 2022-10-23 | Fall | 17.4960 | 0.0603 | 17.6932 | 0.0549 | 0.1973 |
| 2022-10-24 | Fall | 16.1651 | 0.0877 | 16.3979 | 0.0434 | 0.2328 |
| 2022-10-25 | Fall | 21.8243 | 0.1784 | 22.3297 | 0.1515 | 0.5053 |
| 2022-10-26 | Fall | 20.3182 | 0.0662 | 20.5823 | 0.0689 | 0.2640 |
| 2022-10-27 | Fall | 18.1272 | 0.2647 | 19.0792 | 0.5119 | 0.9519 |
| 2022-10-28 | Fall | 15.2825 | 0.1195 | 15.6515 | 0.1310 | 0.3690 |
| 2022-10-29 | Fall | 15.9460 | 0.2919 | 17.0235 | 0.5328 | 1.0776 |
| 2022-10-30 | Fall | 17.8075 | 0.2550 | 18.8869 | 0.4937 | 1.0794 |
| 2022-10-31 | Fall | 18.4347 | 0.0636 | 18.6561 | 0.0643 | 0.2214 |
| 2022-11-01 | Fall | 21.3621 | 0.2237 | 22.2180 | 0.2987 | 0.8559 |
| 2022-11-02 | Fall | 20.7360 | 0.2157 | 21.6695 | 0.3923 | 0.9335 |
| 2022-11-03 | Fall | 20.8683 | 0.2478 | 21.8836 | 0.4639 | 1.0153 |
| 2022-11-04 | Fall | 21.8315 | 0.2208 | 22.7952 | 0.4279 | 0.9637 |
| 2022-11-05 | Fall | 25.2885 | 0.1013 | 25.7796 | 0.1768 | 0.4910 |
| 2022-11-06 | Fall | 23.9980 | 0.0865 | 24.2316 | 0.0703 | 0.2336 |
| 2022-11-07 | Fall | 24.0061 | 0.2928 | 24.5081 | 0.2400 | 0.5019 |
| 2022-11-08 | Fall | 15.1232 | 0.2386 | 15.6634 | 0.2310 | 0.5402 |

**Table S2.**
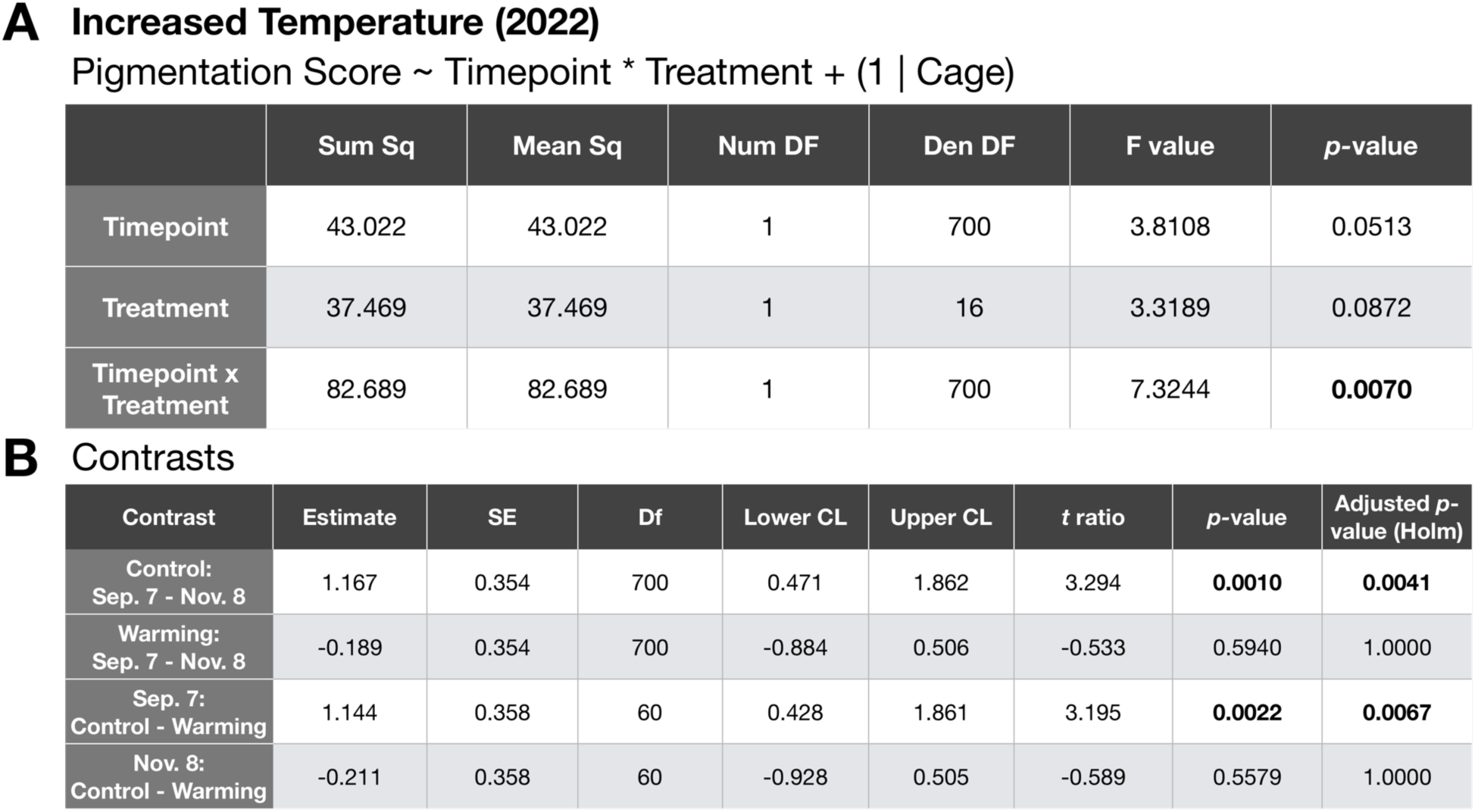
Statistical analyses of the warming treatment. (A) We ran a linear mixed effects model to determine how timepoint, treatment (control vs. warmed temperature), and their interaction influenced pigmentation patterns. Cage was included in the model as a random effect. (B) Planned comparisons showing whether the control and warming treatments exhibited significant shifts in pigmentation from summer to fall, and whether there were significant differences in pigmentation between treatments at the end of the summer (Sep. 7) and fall (Nov. 8) phases. Raw *p-*values and 95% confidence intervals are reported, as well as *p-*values following Holm correction for multiple comparisons.

**Table S3.** Statistical analyses of the reduced intraspecific competition treatment (2017). (A) We ran a linear mixed effects model to determine how timepoint, treatment (control vs. reduced population density), and their interaction influenced pigmentation patterns. Cage was included in the model as a random effect. (B) Planned comparisons showing whether the control and reduced density treatments exhibited significant shifts in pigmentation from summer to fall, and whether there were significant differences in pigmentation between treatments at the end of the summer (Sep. 22) and fall (Nov. 12) phases. Raw *p-*values and 95% confidence intervals are reported, as well as *p-*values following Holm correction for multiple comparisons.

|  | Sum Sq | Mean Sq | Num DF | Den DF | F value | *p*-value |
| --- | --- | --- | --- | --- | --- | --- |
| Timepoint | 1532.6 | 510.87 | 3 | 1079.96 | 23.045 | < .0001 |
| Treatment | 1094.6 | 1094.57 | 1 | 13.88 | 49.375 | < .0001 |
| Timepoint x Treatment | 6271.3 | 2090.42 | 3 | 1079.96 | 94.296 | < .0001 |

B Contrasts
| Contrast | Estimate | SE | Df | Lower CL | Upper CL | *t* ratio | *p*-value | Adjusted *p*-value (Holm) |
| --- | --- | --- | --- | --- | --- | --- | --- | --- |
| Control: Sep. 22 - Nov. 12 | 2.66 | 0.631 | 1082.9 | 1.42 | 3.896 | 4.209 | <.0001 | 0.0001 |
| Reduced Density: Sep. 22 - Nov. 12 | -7.61 | 0.676 | 1081.5 | -8.93 | -6.281 | -11.256 | <.0001 | <.0001 |
| Sep. 22: Control - Density | -2.82 | 1.001 | 48.0 | -4.83 | -0.805 | -2.815 | 0.0071 | 0.0071 |
| Nov. 12: Control - Density | -13.08 | 0.861 | 27.8 | -14.85 | -11.319 | -15.202 | <.0001 | <.0001 |

**Table S4.** Statistical analyses of the reduced intraspecific competition treatment (2022). (A) We ran a linear mixed effects model to determine how timepoint, treatment (control vs. reduced population density), and their interaction influenced pigmentation patterns. Cage was included in the model as a random effect. (B) Planned comparisons showing whether the control and reduced density treatments exhibited significant shifts in pigmentation from summer to fall, and whether there were significant differences in pigmentation between treatments at the end of the summer (Sep. 7) and fall (Nov. 8) phases. Raw *p-*values and 95% confidence intervals are reported, as well as *p-*values following Holm correction for multiple comparisons.

|  | Sum Sq | Mean Sq | Num DF | Den DF | F value | *p*-value |
| --- | --- | --- | --- | --- | --- | --- |
| Timepoint | 7.401 | 7.401 | 1 | 700 | 0.7532 | 0.3858 |
| Treatment | 19.700 | 19.700 | 1 | 16 | 2.0047 | 0.1760 |
| Timepoint x Treatment | 167.235 | 167.235 | 1 | 700 | 17.0182 | < .0001 |

B Contrasts
| Contrast | Estimate | SE | Df | Lower CL | Upper CL | *t* ratio | *p*-value | Adjusted *p*-value (Holm) |
| --- | --- | --- | --- | --- | --- | --- | --- | --- |
| Control: Sep. 7 - Nov. 8 | 1.167 | 0.330 | 700.0 | 0.518 | 1.815 | 3.531 | **0.0004** | **0.0018** |
| Reduced Density: Sep. 7 - Nov. 8 | -0.761 | 0.330 | 700.0 | -1.410 | -0.112 | -2.303 | **0.0216** | **0.0431** |
| Sep. 7: Control - Density | 0.494 | 0.406 | 35.6 | -0.328 | 1.317 | 1.219 | 0.2309 | 0.2309 |
| Nov. 8: Control - Density | -1.433 | 0.406 | 35.6 | -2.256 | -0.610 | -3.534 | **0.0012** | **0.0035** |

**Table S5.** Statistical analyses of the interspecific competition treatment. (A) We ran a linear mixed effects model to determine how timepoint, treatment (control vs. competition with *Z. indianus*), and their interaction influenced pigmentation patterns. Cage was included in the model as a random effect. (B) Planned comparisons showing whether the control and competition treatments exhibited significant shifts in pigmentation from summer to fall, and whether there were significant differences in pigmentation between treatments at the end of the summer (Sep. 11) and fall (Nov. 8) phases. Raw *p-*values and 95% confidence intervals are reported, as well as *p-*values following Holm correction for multiple comparisons.

|  | Sum Sq | Mean Sq | Num DF | Den DF | F value | *p*-value |
| --- | --- | --- | --- | --- | --- | --- |
| Timepoint | 332.56 | 332.56 | 1 | 505 | 23.0331 | < .0001 |
| Treatment | 22.30 | 22.30 | 1 | 11 | 1.5445 | 0.2398 |
| Timepoint x Treatment | 32.85 | 32.85 | 1 | 505 | 2.2751 | 0.1321 |

B Contrasts
| Contrast | Estimate | SE | Df | Lower CL | Upper CL | *t* ratio | *p*-value | Adjusted *p*-value (Holm) |
| --- | --- | --- | --- | --- | --- | --- | --- | --- |
| Control:<br>Sep. 11 - Nov. 8 | 1.1000 | 0.454 | 505.0 | 0.208 | 1.9923 | 2.422 | **0.0158** | **0.0474** |
| Competition:<br>Sep. 11 - Nov. 8 | 2.1083 | 0.491 | 505.0 | 1.145 | 3.0721 | 4.298 | **<.0001** | **0.0001** |
| Sep. 11:<br>Control - Competition | -1.0512 | 0.553 | 27.1 | -2.185 | 0.0825 | -1.902 | 0.0678 | 0.1357 |
| Nov. 8:<br>Control - Competition | -0.0429 | 0.553 | 27.1 | -1.177 | 1.0909 | -0.078 | 0.9388 | 0.9388 |

**Table S6.** Statistical analyses of the diet treatment. (A) We ran a linear mixed effects model to determine how timepoint, treatment (control vs. apple food), and their interaction influenced pigmentation patterns. Cage was included in the model as a random effect. (B) Planned comparisons showing whether the control and apple treatments exhibited significant shifts in pigmentation from summer to fall, and whether there were significant differences in pigmentation between treatments at the end of the summer (Sep. 7) and fall (Nov. 24) phases. Raw *p-*values and 95% confidence intervals are reported, as well as *p-*values following Holm correction for multiple comparisons.

|  | Sum Sq | Mean Sq | Num DF | Den DF | F value | *p*-value |
| --- | --- | --- | --- | --- | --- | --- |
| Timepoint | 428.36 | 107.089 | 4 | 880 | 10.1293 | < .0001 |
| Treatment | 19.12 | 19.123 | 1 | 10 | 1.8088 | 0.2084 |
| Timepoint x Treatment | 175.07 | 43.768 | 4 | 880 | 4.1399 | 0.0025 |

**B Contrasts**
| Contrast | Estimate | SE | Df | Lower CL | Upper CL | *t* ratio | *p*-value | Adjusted *p*-value (Holm) |
| --- | --- | --- | --- | --- | --- | --- | --- | --- |
| Control:<br>Sep. 7 - Nov. 24 | 1.63 | 0.485 | 880 | 0.682 | 2.585 | 3.370 | 0.0008 | 0.0024 |
| Apple:<br>Sep. 7 - Nov. 24 | 1.72 | 0.485 | 880 | 0.771 | 2.674 | 3.553 | 0.0004 | 0.0016 |
| Sep. 7:<br>Control - Apple | -1.23 | 0.496 | 160 | -2.213 | -0.254 | -2.486 | 0.0139 | 0.0279 |
| Nov. 24:<br>Control - Apple | -1.14 | 0.496 | 160 | -2.124 | -0.165 | -2.307 | 0.0224 | 0.0279 |

**Table S7.** Statistical analyses of the *A. thailandicus* treatment. We manipulated the gut microbiome by supplementing apple food with *Acetobacter thailandicus*; the control in this experiment was apple food. (A) We ran a linear mixed effects model to determine how timepoint, treatment (control vs. *A. thailandicus*), and their interaction influenced pigmentation patterns. Cage was included in the model as a random effect. (B) Planned comparisons showing whether the control and *A. thailandicus* treatments exhibited significant shifts in pigmentation from summer to fall, and whether there were significant differences in pigmentation between treatments at the end of the summer (Sep. 7) and fall (Nov. 24) phases. Raw *p-*values and 95% confidence intervals are reported, as well as *p-*values following Holm correction for multiple comparisons.

|  | Sum Sq | Mean Sq | Num DF | Den DF | F value | *p*-value |
| --- | --- | --- | --- | --- | --- | --- |
| Timepoint | 174.448 | 87.224 | 2 | 524 | 7.8460 | **0.0004** |
| Treatment | 4.859 | 4.859 | 1 | 10 | 0.4371 | 0.5235 |
| Timepoint x Treatment | 105.100 | 52.550 | 2 | 524 | 4.7270 | **0.0092** |

**B Contrasts**
| Contrast | Estimate | SE | Df | Lower CL | Upper CL | *t* ratio | *p*-value | Adjusted *p*-value (Holm) |
| --- | --- | --- | --- | --- | --- | --- | --- | --- |
| Control (Apple):<br>Sep. 7 - Nov. 24 | 1.722 | 0.497 | 524 | 0.7458 | 2.70 | 3.465 | **0.0006** | **0.0023** |
| At (Apple + At):<br>Sep. 7 - Nov. 24 | 0.956 | 0.497 | 524 | -0.0209 | 1.93 | 1.923 | 0.0551 | 0.1102 |
| Sep. 7:<br>Control - At | 1.189 | 0.527 | 58 | 0.1341 | 2.24 | 2.256 | **0.0278** | 0.0835 |
| Nov. 24:<br>Control - At | 0.422 | 0.527 | 58 | -0.6326 | 1.48 | 0.801 | 0.4263 | 0.4263 |

**Table S8.** Statistical analyses of the *L. brevis* treatment. We manipulated the gut microbiome by supplementing apple food with *Lactobacillus brevis*; the control in this experiment was apple food. (A) We ran a linear mixed effects model to determine how timepoint, treatment (control vs. *L. brevis*), and their interaction influenced pigmentation patterns. Cage was included in the model as a random effect. (B) Planned comparisons showing whether the control and *L. brevis* treatments exhibited significant shifts in pigmentation from summer to fall, and whether there were significant differences in pigmentation between treatments at the end of the summer (Sep. 7) and fall (Nov. 24) phases. Raw *p-*values and 95% confidence intervals are reported, as well as *p-*values following Holm correction for multiple comparisons.

|  | Sum Sq | Mean Sq | Num DF | Den DF | F value | *p*-value |
| --- | --- | --- | --- | --- | --- | --- |
| Timepoint | 70.415 | 35.207 | 2 | 524 | 3.4758 | **0.0317** |
| Treatment | 13.399 | 13.399 | 1 | 10 | 1.3227 | 0.2769 |
| Timepoint x Treatment | 169.615 | 84.807 | 2 | 524 | 8.3724 | **0.0003** |

**B Contrasts**
| Contrast | Estimate | SE | Df | Lower CL | Upper CL | *t* ratio | *p*-value | Adjusted *p*-value (Holm) |
| --- | --- | --- | --- | --- | --- | --- | --- | --- |
| Control (Apple):<br>Sep. 7 - Nov. 24 | 1.722 | 0.474 | 524.0 | 0.790 | 2.654 | 3.630 | **0.0003** | **0.0012** |
| *Lb* (Apple + *Lb*):<br>Sep. 7 - Nov. 24 | -0.656 | 0.474 | 524.0 | -1.588 | 0.276 | -1.382 | 0.1676 | 0.1676 |
| Sep. 7:<br>Control - *Lb* | 1.244 | 0.488 | 69.5 | 0.272 | 2.217 | 2.552 | **0.0129** | **0.0388** |
| Nov. 24:<br>Control - *Lb* | -1.133 | 0.488 | 69.5 | -2.106 | -0.161 | -2.324 | **0.0231** | **0.0461** |

